# Plastidial (p)ppGpp synthesis by the Ca^2+^-dependent RelA-SpoT homolog regulates the adaptation of chloroplast gene expression to darkness in *Arabidopsis*

**DOI:** 10.1101/767004

**Authors:** Sumire Ono, Sae Suzuki, Doshun Ito, Shota Tagawa, Takashi Shiina, Shinji Masuda

**Author notes:** These authors contributed equally to the study (Sumire Ono, Sae Suzuki).

## Abstract

In bacteria, the hyper-phosphorylated nucleotides, guanosine 5’-diphosphate 3’-diphosphate (ppGpp) and guanosine 5’-triphosphate 3’-diphosphate (pppGpp), function as secondary messengers in the regulation of various metabolic processes of the cell, including transcription, translation, and enzymatic activities, especially under nutrient deficiency. The activity carried out by these nucleotide messengers is known as the stringent response. (p)ppGpp levels are controlled by two distinct enzymes, namely, RelA and SpoT, in *Escherichia coli*. RelA-SpoT homologs (RSHs) are also conserved in plants and algae where they function in the plastids. The model plant *Arabidopsis thaliana* contains four RSHs: RSH1, RSH2, RSH3, and Ca^2+^-dependent RSH (CRSH). Genetic characterizations of RSH1, RSH2, and RSH3 were undertaken, which showed that the (p)ppGpp-dependent plastidial stringent response significantly influences plant growth and stress acclimation. However, the physiological significance of CRSH-dependent (p)ppGpp synthesis remains unclear, as no *crsh*-null mutant has been available. Here to investigate the function of CRSH, a *crsh*-knockout mutant of *Arabidopsis* was constructed using a site-specific gene-editing technique, and its phenotype was characterized. A transient increase of ppGpp was observed for 30 min in the wild type (WT) after light-to-dark transition, but this increase was not observed in the *crsh* mutant. Similar analyzes were performed with the *rsh2rsh3* double and *rsh1rsh2rsh3* triple mutants of *Arabidopsis* and showed that the transient increments of ppGpp in the mutants were higher than those in the WT. The increase of (p)ppGpp in the WT and *rsh2rsh3* accompanied decrements in the mRNA levels of *psbD* transcribed by the plastid-encoded plastid RNA polymerase. These results indicated that the transient increase of intracellular ppGpp at night is due to CRSH-dependent ppGpp synthesis and the (p)ppGpp level is maintained by the hydrolytic activities of RSH1, RSH2, and RSH3 to accustom plastidial gene expression to darkness.

## Introduction

Organisms have various mechanisms by which they adapt their physiology to environmental changes. Autophagy is a famous nutritional starvation response in eukaryotes (Nakatogawa and Ohsumi 2014; Levine and Klionsky 2017), but this process does not occur in bacteria. Instead, the well-known stringent response provides the bacterial reaction to nutrient starvation (Potrykus and Cashel 2008; Dalebroux and Swanson 2012). The stringent response was discovered almost a half-century ago as a reaction that reduces the synthesis of rRNA and tRNA when *Escherichia coli* is exposed to amino-acid starvation (Cashel 1969). This response is controlled by the hyper-phosphorylated nucleotides guanosine 5’-diphosphate 3’-diphosphate (ppGpp) and guanosine 5’-triphosphate 3’-diphosphate (pppGpp). Because these molecules are similar in function, they are collectively called (p)ppGpp. Besides nutrient starvation, the (p)ppGpp-dependent stringent response is important to combating multiple stresses, including fatty-acid deficiency and high temperatures, and is conserved among almost all bacteria (Potrykus and Cashel 2008).

The stringent response regulates various metabolic processes, including transcription, translation, and enzymatic activities (Potrykus and Cashel 2008). In *E. coli*, (p)ppGpp is synthesized by two enzymes, namely, RelA and SpoT, whereas degradation is performed by SpoT. Thus, SpoT catalyzes (p)ppGpp synthesis and hydrolysis, whereas RelA functions as a (p)ppGpp synthase (Fig. 1). The stringent response has been studied extensively in bacteria, but in recent years, RelA-SpoT homologs (RSHs) have been discovered in plants and algae (Tozawa and Nomura 2011; Masuda 2012; Boniecka et al. 2017; Field 2018). RSHs were introduced into an ancestral proto-plant cell through multiple horizontal gene transfers from two distinct bacterial phyla, including *Deinococcus–Thermus* (Ito et al. 2017). *Arabidopsis thaliana* contains four nuclear-encoded RSHs: RSH1, RSH2, RSH3, and Ca^2+^-dependent RSH (CRSH) (Fig. 1). RSH1 may only have (p)ppGpp hydrolase activity because it lacks an amino-acid necessary for (p)ppGpp synthase activity. RSH2 and RSH3 show high similarity (~80% amino-acid identity) and may be bifunctional enzymes with (p)ppGpp synthase and hydrolase activities. CRSH lacks a histidine-aspartate (HD) domain necessary for (p)ppGpp hydrolase activity and functions only as a (p)ppGpp synthase (Ito et al. 2017). CRSH has two Ca^2+^-binding EF hand motifs at the C-terminus, and its (p)ppGpp synthase activity requires Ca^2+^ *in vitro* (Tozawa et al. 2007). Thus, *Arabidopsis* RSHs have been classified into three distinct groups, namely, RSH1, RSH2/3, and CRSH, all of which are localized in plastids. These classes of RSHs are well conserved in many plant and algal species (Ito et al. 2017), suggesting that (p)ppGpp levels in plant cells are maintained by coordinated regulation of their enzyme activities.

**Fig. 1.**
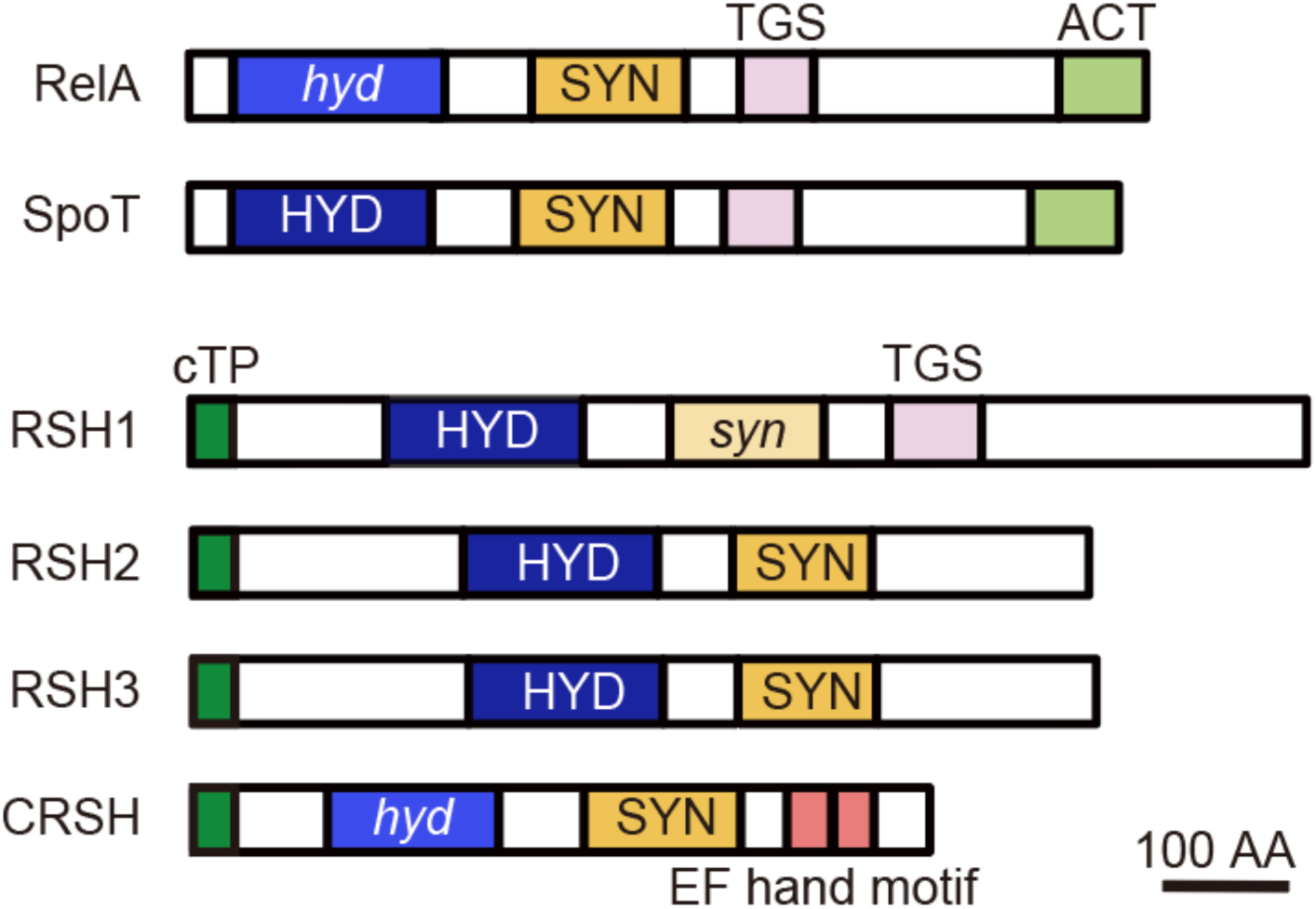
Domain structures of RelA and SpoT from *E. coli* and RSHs in *Arabidopsis*. Hyd, (p)ppGpp hydrolase domain; *hyd*, (p)ppGpp hydrolase domain lacking some critical amino acids; Syn, (p)ppGpp synthase domain; *syn*, (p)ppGpp synthase domain lacking the critical Gly residue; ACT, aspartate kinase, chorismate mutase, TyrA domain; EF hand, Ca^2+^-binding helix E and F motif; TGS, threonyl-tRNA synthase, GTPase SpoT domain; AA, amino acids; cTP, chloroplast transit peptide.

The effects of (p)ppGpp on plant growth have been investigated using *RSH3*-overexpression lines that over-accumulate ppGpp (Maekawa et al. 2015; Sugliani et al. 2016; Honoki et al. 2018). Mutant-specific phenotypes include pale-green leaves, dwarfization of chloroplasts, increased tolerance to nitrogen deficiency, and reduced mRNA levels of chloroplast genes, which indicate that the plastidial stringent response affects plant growth and adaptation to stresses, although the exact targets of (p)ppGpp in chloroplasts are largely unclarified. The artificial accumulation of (p)ppGpp in the cytosol induces a dwarf phenotype in *Arabidopsis* (Ihara and Masuda 2016), which indicates that the (p)ppGpp-dependent stringent response may function in the cytosol; however, this hypothesis requires confirmation.

In plants, Ca^2+^ functions as a major secondary messenger that is triggered by various stresses, including wounding, darkness, and low temperatures (Knight et al. 1996; Sanders et al. 1999; Nguyen et al. 2018). A large fraction of Ca^2+^ is stored in the vacuoles and endoplasmic reticulum of plant cells (White and Broadley 2003). In addition, chloroplasts and mitochondria are used as Ca^2+^ storage sites and/or sinks (Sanders et al. 1999). Plastidial Ca^2+^ concentrations alter in response to environmental conditions (Hochmal et al. 2015). When plants are placed in the dark, the Ca^2+^ concentration in the chloroplast stroma transiently increases (Sai and Johnson 2002; Loro et al. 2016), which is thought to occur due to the release of Ca^2+^ from the thylakoid membranes and/or its uptake across the chloroplast envelope by envelope-localized proteins that include BICAT2 (Hochmal et al. 2015; Frank et al. 2019). The significance of the increase in Ca^2+^ content upon the transition to darkness has been long discussed. A recent prediction is that Ca^2+^ plays a role in shutting down unnecessary photosynthetic reactions at night by suppressing enzymatic activities involved in the Calvin–Benson cycle (Portis and Heldt 1976; Charles and Halliwell 1980; Hertig and Wolosiuk 1980; Rocha et al. 2014). The *Arabidopsis bicat2* mutation, which displays low Ca^2+^ uptake activity under transition to darkness, causes significant growth retardation. The *bicat2* phenotype can be restored in part by the addition of sucrose (Frank et al. 2019), which may indicate that the negative impact of the *bicat2* mutation on plant growth is due to an imbalance in the control of photosynthesis. Therefore, Ca^2+^-dependent signaling in chloroplasts is important for the adaptation of photosynthetic activity to darkness, although the mechanisms are not fully understood.

Plastidial Ca^2+^ signaling also triggers protective responses against pathogens. When plants encounter infectious agents, such as pathogenic bacteria, the plastidial Ca^2+^ concentration increases, which further signals the nucleus to induce the transcription of genes related to the defense response, including salicylic acid biosynthesis (Nomura et al. 2012). Specifically, the mutational loss of the plastidial protein, namely, CAS, results in the inability to increase Ca^2+^ concentrations in chloroplasts upon pathogen attack and the incapacity to upregulate nuclear gene expression. However, the mechanisms underlying the plastidial Ca^2+^-based retrograde signal to the nucleus remain largely unknown.

A potential relationship between plastidial Ca^2+^ signaling and the (p)ppGpp-dependent stringent response has been suggested, as the (p)ppGpp synthase activity of CRSH requires Ca^2+^ *in vitro* (Tozawa et al. 2007). However, this hypothesis remains to be tested, as no *crsh*-null mutant has been available. Here, to understand the physiological significance of CRSH function, we constructed an *Arabidopsis crsh*-null mutant using the CRISPR/Cas9-based gene-editing technique and characterized the mutant phenotype. The results indicate that the darkness-induced accumulation of (p)ppGpp relies on CRSH activity, which is necessary to downregulate some plastidial gene expression and adapt the photosynthetic process to dark conditions.

## Results

### Isolation of the *crsh*-null mutant

Previously, a T-DNA insertional *Arabidopsis crsh* mutant *crsh-1* (SAIL_1295_C04) was reported to have a T-DNA insertion in the 3’-UTR of *CRSH* (Sugliani et al. 2016). We obtained this mutant line from the stock center and the homozygous mutant was isolated via PCR-based genotyping (Fig. 2B). Contrary to the previous report, our sequencing analysis indicated that *crsh-1* actually has the T-DNA insertion in the *CRSH* coding region (Fig. 2A), suggesting that the T-DNA insertion may inactivate *CRSH*. However, the T-DNA insertion creates a stop codon just after the T-DNA insertion site, and the deduced molecular weight of the mutated CRSH is similar to (~0.7 kDa less than) that of the wild type (WT). Reverse-transcription (RT)-PCR and western blot analyzes clearly showed mRNA and protein accumulation, respectively, from the mutated CRSH in *crsh-1* (Figs. 2C, D), suggesting that the mutated CRSH is functional in *crsh-1*. We therefore assumed that the isolation of the *crsh*-null mutant using other strategies was required to further characterize CRSH function.

**Fig. 2.**
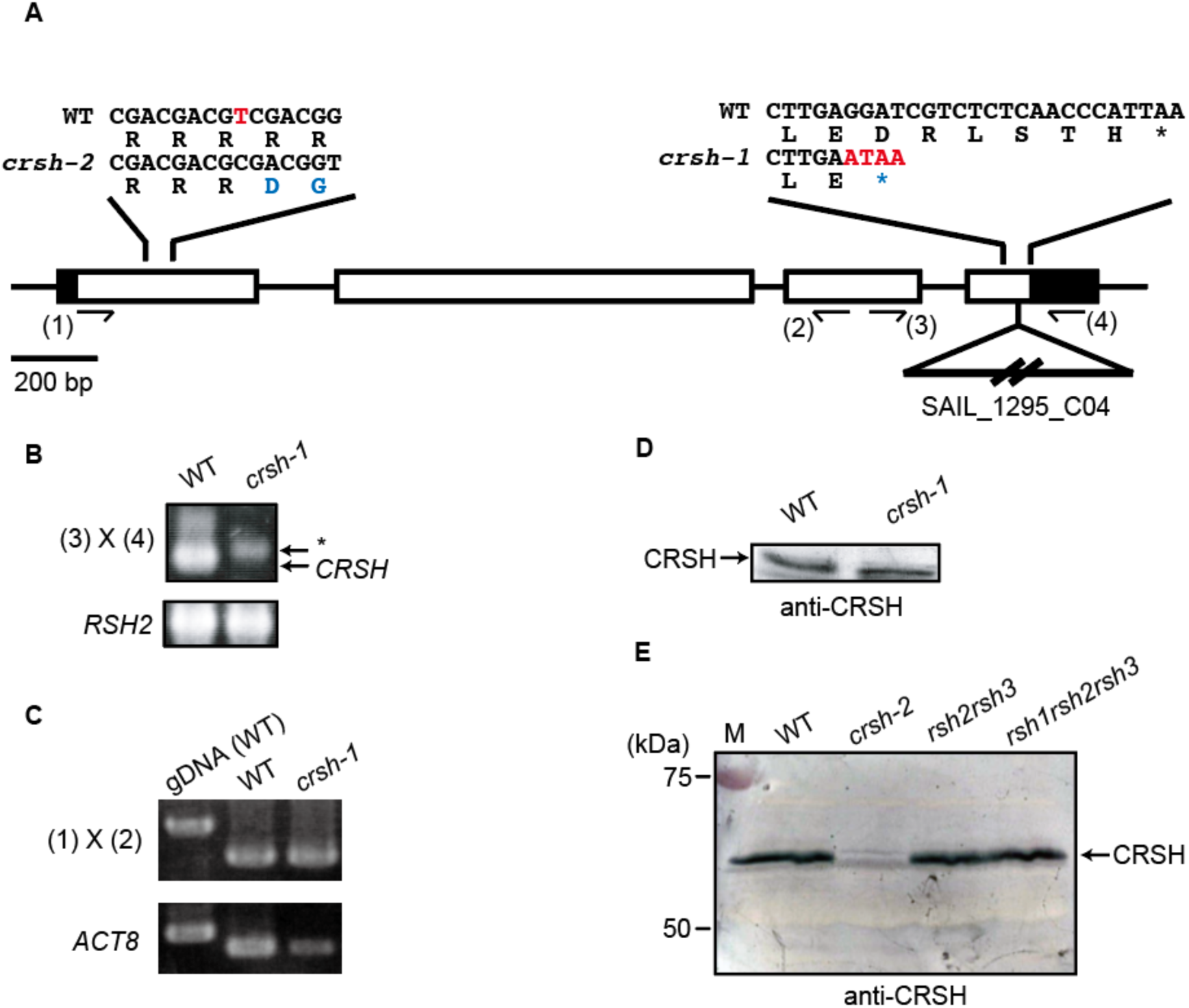
Construction of *crsh* mutants. (A) Schematic of the exon–intron structure of *CRSH* (At3g17470) with a single nucleotide deletion site in *crsh-2* and a T-DNA insertion site in *crsh-1*. A deleted nucleotide in *crsh-2* and inserted T-DNA nucleotides in *crsh-1* are highlighted in red. Mutated codons are highlighted in blue. (B) PCR analysis reveals homozygous T-DNA insertion in *crsh-1*. Positions of primers used (3 and 4) are indicated in (A). An asterisk indicates a nonspecific band. *RSH2* was amplified as a control for DNA isolation. (C) RT-PCR analysis reveals the presence of *CRSH* mRNA in *crsh-1*. Positions of primers used (1 and 2) are indicated in (A). Genome DNA isolated from WT was used as a control for genome DNA contamination, and the *actin8* (ACT8) gene was used as a control for mRNA isolation and reverse transcription. (D) Western blot of proteins isolated from WT and *crsh-1*. (E) Western blot of proteins isolated from WT, *crsh-2, rsh2rsh3*, and *rsh1rsh2rsh3*. Full images of original photographs and western blot data are shown in Supplementary Fig. S1.

We employed a CRISPR/CAS9-based gene-editing method (Fauser et al. 2014) to obtain the *crsh*-null mutant, and designed a protospacer sequence in the first exon of *CRSH* to induce a double-strand break at the correct position. Approximately 20 independent T1 plants were isolated, and one mutant showing a single base deletion at the 99th nucleotide in the first exon of *CRSH* was found in the T3 generation (Fig. 2A).

This mutation induced a frameshift, which created a stop codon at the 72nd codon in the first exon (the 38th codon from the frameshift site). Western blot analysis using an anti-CRSH antibody confirmed the absence of CRSH in the mutant (Fig. 2E); we designated the *crsh*-null mutant as *crsh-2*.

### ppGpp accumulation in *rsh* mutants

Subsequently, we quantified ppGpp levels in *crsh-2* to evaluate the contribution of CRSH to ppGpp synthesis. No significant difference was observed in ppGpp levels between the WT (172.9 ± 15.6 pmol g-1) and *crsh-2* (186.8 ± 5.6 pmol g-1) under continuous light conditions, indicating that CRSH is not required to synthesize basal levels of ppGpp. Upon light-to-dark transition, the WT exhibited an increase in ppGpp for 30 min, which decreased to basal levels after 60 min (Fig. 3A). The dark-induced transient increment of ppGpp was also observed in *crsh-2* after 10 min; however, the increment rapidly decreased to low levels after 30 min (Fig. 3A). A complementing line of *crsh-2*, harboring a 35S-promoter-driven *CRSH*, maintained high levels of ppGpp beyond 30 min after the transition to darkness (Supplementary Fig. S2). This result indicates that the attenuation of ppGpp levels upon dark treatment is due to the upregulation of CRSH enzyme activity. Further support for this hypothesis was established by measuring ppGpp levels following light-to-dark transition in the *rsh2rsh3* double mutant. Besides CRSH, the *rsh2rsh3* mutant contains no ppGpp synthase, as RSH1 has (p)ppGpp hydrolase activity but no (p)ppGpp synthase activity.

**Fig. 3.**
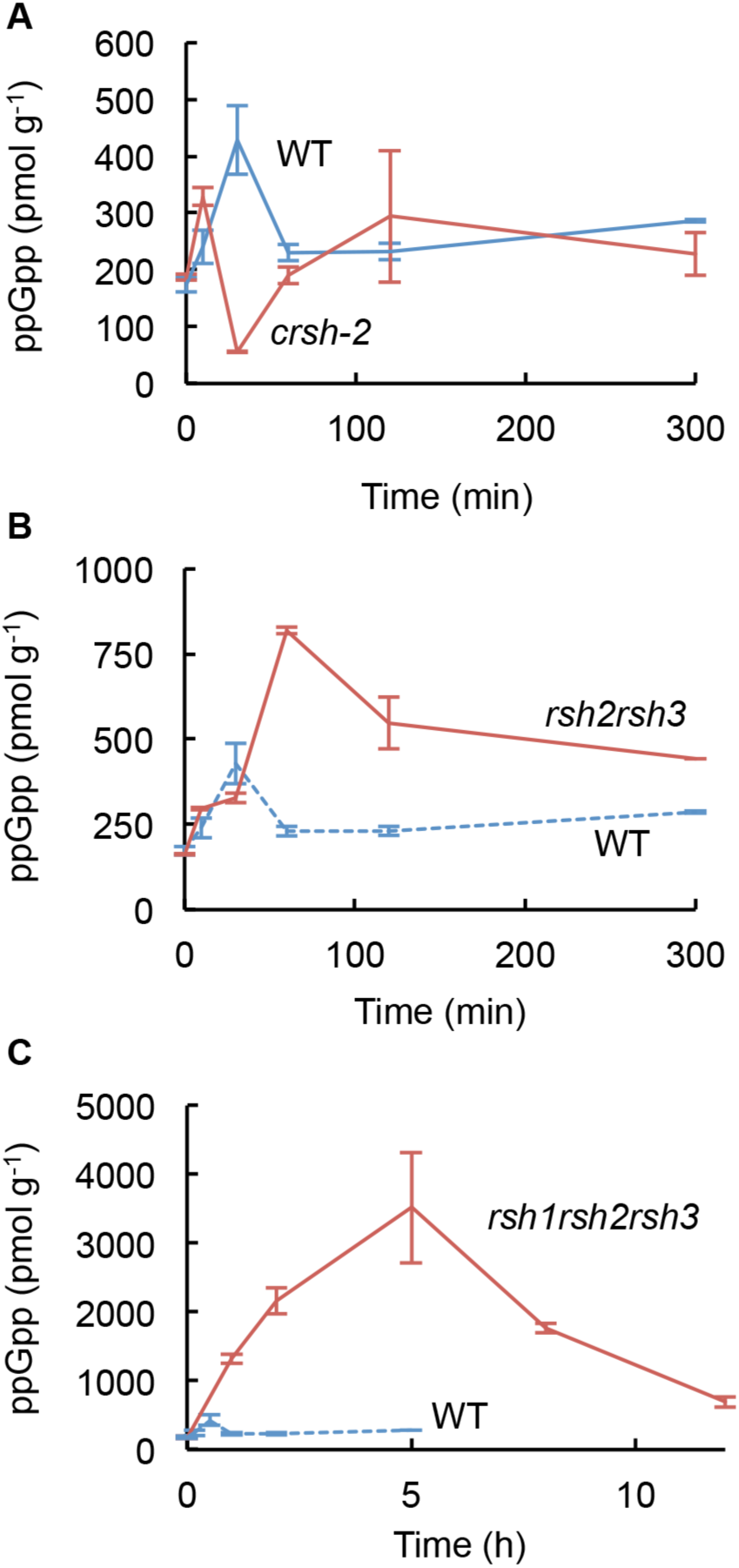
ppGpp accumulation kinetics upon dark-to-light transition in the WT and *crsh-2* (A), WT and *rsh2rsh3* (B), and WT and *rsh1rsh2rsh3* (C). Plants were grown on 0.8% agar-solidified 1/2 MS medium under continuous light (40 µmol photons m^−2^ s^−1^). Fourteen-day-old plants were transferred to the dark and shoots were harvested at the time points for ppGpp quantification. Values are mean ± SD (*n* = 3). Data for the WT are reproduced (dotted lines).

As reported previously (Maekawa et al. 2015), the amount of ppGpp in *rsh2rsh3* before the light-to-dark transition (162.5 ± 3.0 pmol g^−1^) was similar to that in the WT (as indicated above). Upon light-to-dark transition, *rsh2rsh3* exhibited a continuously raised concentration of ppGpp for 60 min (Fig. 3B). Peak ppGpp levels in the WT and *rsh2rsh3* were 428 ± 60 pmol g^−1^ at 30 min and 818 ± 10 pmol g^−1^ at 60 min after the transition to darkness, respectively. CRSH functions as the sole (p)ppGpp synthase in the *rsh2rsh3* mutant, indicating that increases in (p)ppGpp upon the shift from light-to-dark are due to the upregulation of CRSH activity.

We also measured the ppGpp kinetics in the *rsh1rsh2rsh3* triple mutant, which lacks all (p)ppGpp-specific hydrolases. Before dark treatment, no significant difference in ppGpp levels was observed between *rsh1rsh2rsh3* (176.0 ± 18.5 pmol g-1) and the WT (as indicated above). Following light-to-dark transition, ppGpp levels were significantly raised for ~5 h in the *rsh1rsh2rsh3* mutant (Fig. 3C). The ppGpp concentration 5 h after the transition to darkness was 3,517 ± 807 pmol g^−1^, which was ~10- and ~5-fold higher than those found in the WT (after 30 min) and *rsh2rsh3* mutant (~2 h), respectively. These results indicated that CRSH is responsible for the upregulation of ppGpp synthesis upon light-to-dark transition, and the hydrolase activities of RSH1, RSH2, and RSH3 are required to maintain adequate levels of (p)ppGpp during the night.

Western blot analysis using the anti-CRSH antibody was performed to establish the CRSH levels in the *rsh2rsh3* and *rsh1rsh2rsh3* mutants. No differences in CRSH levels were found in either mutant (Fig. 2E), indicating that the accumulation of ppGpp upon transition to darkness in the mutants was not due to a change in CRSH levels.

### Dark-induced ppGpp accumulation regulates the transcription of chloroplast genes

To investigate the effects of darkness-induced increases in ppGpp on chloroplast gene expression, some chloroplast gene transcripts were measured by quantitative RT (qRT)-PCR from the WT, *crsh-2*, and *rsh2rsh3*. We monitored the mRNA levels of *accD, clpP, atpB, rbcL*, and *psbD*. The *accD* and *clpP* genes are transcribed by the nuclear-encoded plastid RNA polymerase (NEP), *rbcL* and *psbD* are transcribed by the plastid-encoded plastid RNA polymerase (PEP), and *atpB* is transcribed by the NEP and PEP. The transcripts of *psbD* in the WT and *rsh2rsh3* were reduced 1 h after light-to-dark transition (Fig. 4). However, no significant reduction was observed in *crsh-2*, suggesting that the reduction of *psbD* mRNA levels was due to CRSH-dependent ppGpp synthesis. A significant reduction of *atpB* mRNA levels was observed in *rsh2rsh3* and the transcript of *clpP* in this mutant rose specifically at 0.5 h from light-to-dark transition (Fig. 4), although the transient increment was not observed in the WT or *crsh-2*. The *accD* and *rbcL* levels were insignificantly altered in any of the tested lines.

**Fig. 4.**
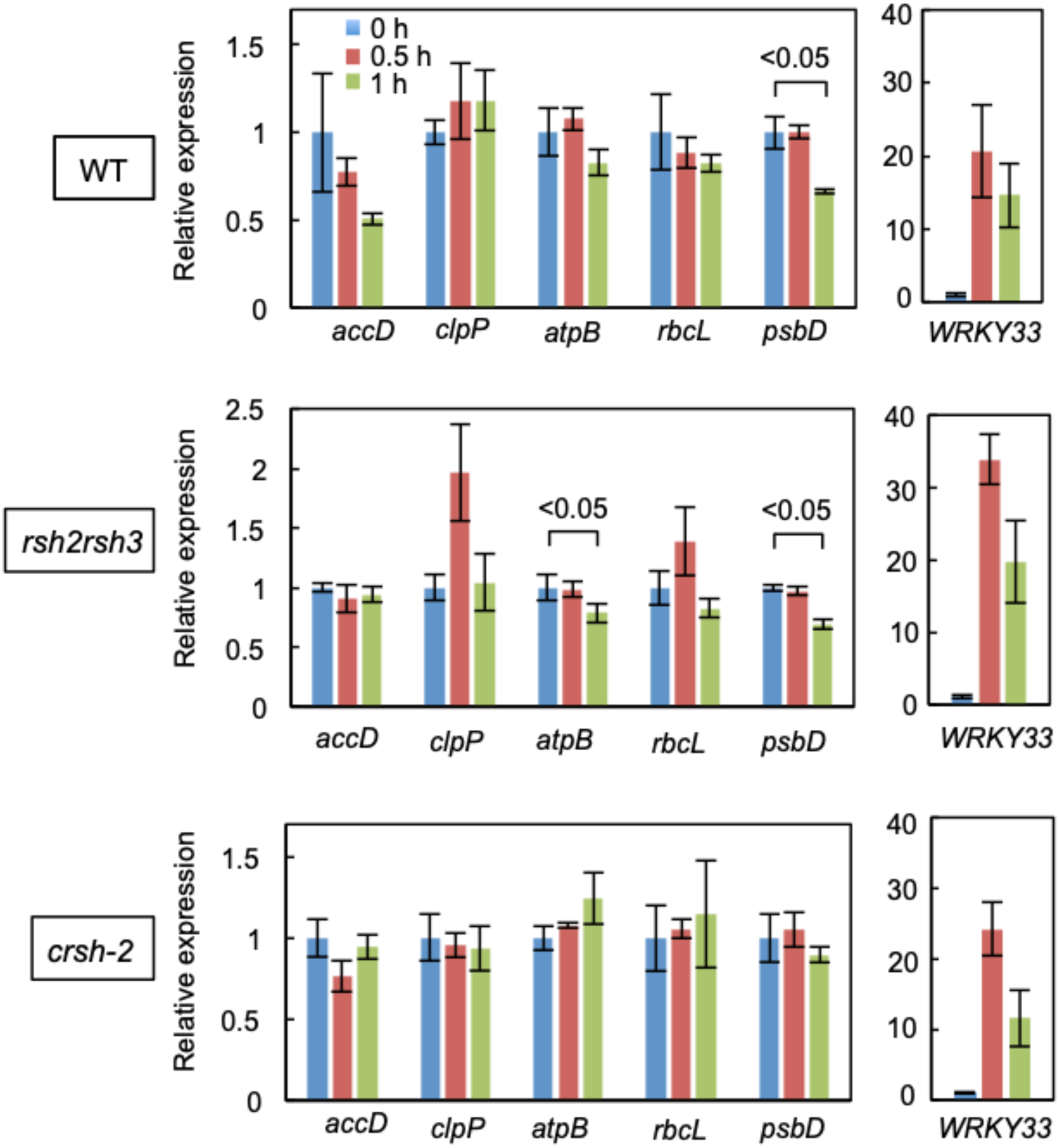
Relative mRNA levels in the WT, *rsh2rsh3*, and *crsh-2* plants before (0 h) and after dark incubation for 0.5 and 1 h. Plants were grown on 0.8% agar-solidified 1/2 MS medium under continuous light (40 µmol photons m^−2^ s^−1^) for 14 days and then transferred to the dark. Messenger RNA quantification was carried out using RT-PCR analysis. Values are mean ± SD (*n* = 3). The WT level at 0 h in the first experiment was set to 1.0. *P* values based on Student’s *t*-test are indicated.

### Transcription profile following pathogen stimulation in the *crsh* mutant

Previous studies have indicated that after the application of pathogen-associated molecular patterns (PAMPs), the Ca^2+^ concentration in the chloroplast stroma increases, and the expression of certain nuclear genes is upregulated as part of the defense response (Nomura and Shiina 2014; Sano et al. 2014). To test the hypothesis that Ca^2+^-dependent (p)ppGpp synthesis is involved in the defense response, we analyzed the mRNA levels of nuclear-encoded *WRKY33* and *WRKY46* upon the addition of a PAMP, namely, flg22, in the WT and *crsh-2* mutant. WRKY33 and WRKY46 were previously shown to be upregulated at the transcriptional level by PAMPs (Sano et al. 2014). As shown in Fig. 5, no significant changes were observed in the transcription profiles for these genes in the WT and *crsh-2*, suggesting that CRSH is not required for the Ca^2+^-dependent signaling associated with exposure to PAMPs.

**Fig. 5.**
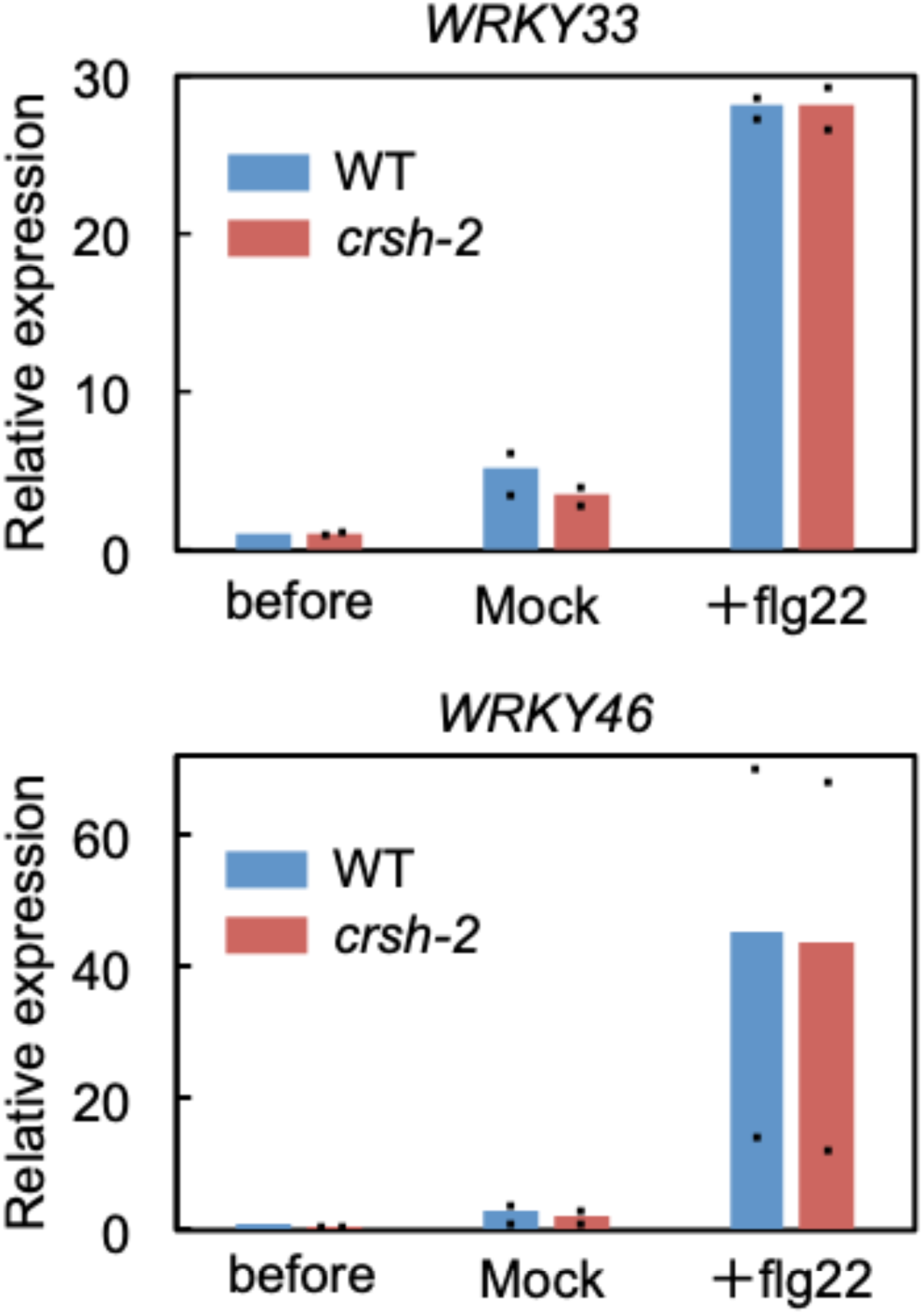
Relative mRNA levels in the WT and *crsh-2* before and after incubation with flg22. Plants were grown on agar-solidified 1/2 MS medium under 16 h light (100 µmol photons m^−2^ s^−1^)/16 h dark cycles. Fourteen-day-old plants were treated with 1 µM flg22 and/or solvent (water) for 0.5 h. Values are means of two independent experiments and the data are indicated by dots. The WT level before flg22 addition was set to 1.0.

To obtain further insight into the role of the plastidial stringent response in plant defense, we analyzed the transcript levels of *WRKY33* upon transition to darkness (Fig. 4). In fact, WRKY33 transcripts accrued significantly after the light-to-dark transition; in the WT, ~20- and ~15-fold higher accumulations of mRNA were observed at 0.5 and 1 h, respectively. A similar expression pattern was observed in *crsh-2*, indicating that CRSH is not responsible for the darkness-induced accumulation of WRKY33 transcripts. However, we found that *rsh2rsh3* showed higher (1.5-fold) levels of *WRKY33* mRNA accumulation than the WT after 0.5 h of dark treatment, suggesting that the ppGpp-dependent plastidial stringent response is involved in plant defense.

### Growth phenotype of *rsh* mutants

To observe the effects of nighttime ppGpp accumulation on plant growth, we grew WT, *crsh-2*, and *rsh1rsh2rsh3* mutants under continuous light, short-day (8 h light/16 h dark) and long-day (16 h light/8 h dark) conditions for 14 days. The fresh weights of WT and *crsh-2* shoots were similar; however, a significant reduction in the fresh weight of *rsh1rsh2rsh3* compared with that of the WT was observed only under short-day conditions (Fig. 6), suggesting that the hyper-accumulation of ppGpp in the triple mutant under dark conditions (Fig. 3C) causes growth retardation. Under long-day conditions, a significant reduction in the fresh weight of *rsh1rsh2rsh3* compared with *crsh-2* was observed (Fig. 6), which supported the hypothesis that dark-induced synthesis of ppGpp downregulates plant growth. Notably, the growth-retarding phenotype of *rsh1rsh2rsh3* was rescued by the exogenous addition of sucrose.

**Fig. 6.**
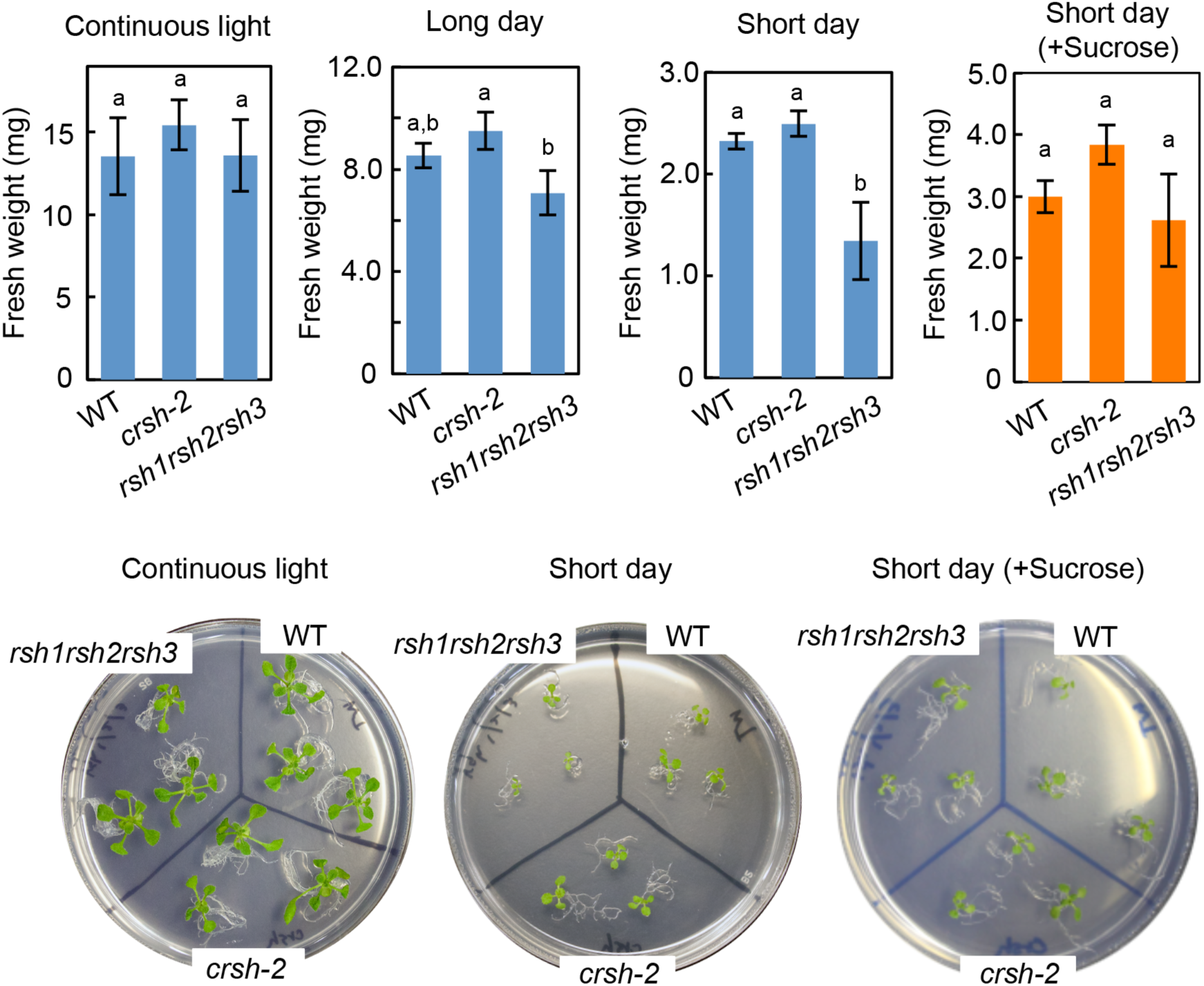
Growth phenotype of *rsh* mutants. Plants were grown on 0.8% agar-solidified 1/2 MS medium with or without 1% sucrose under continuous light, long-day (16 h light/8 h dark) or short-day (8 h light/16 h dark) conditions for 14 days, and then fresh weights of shoots were measured (*n* = 29–46). Different letters denote the significant difference by using Tukey–Kramer’s multiple comparison test (*P <* 0.05).

Previously, an *Arabidopsis crsh* co-suppression mutant (OX19) obtained from 35S-promoter-driven *CRSH*-overexpression lines showed abnormal flower development (Masuda et al. 2008); however, this phenotype was not observed in *crsh-2*. A *CRSH* knockdown mutant constructed using an artificial microRNA also did not show this abnormal phenotype (Sugliani et al. 2016), suggesting that the OX19 phenotype was not simply due to the less accumulation of CRSH. No further morphological differences were observed for *crsh-2* and *rsh2rsh3* in comparison with the WT throughout the entire growth period (Supplementary Fig. S3).

## Discussion

In this study, we provide the first *in vivo* evidence demonstrating the Ca^2+^-dependent ppGpp synthase activity of CRSH. We succeeded in isolating the *Arabidopsis crsh*-null mutant, *crsh-2*, which did not attenuate a transient increase of ppGpp upon shifting from light-to-dark conditions. Given that plastidial Ca^2+^ concentrations transiently increase upon light-to-dark transition (Sai and Johnson 2002), CRSH most likely senses the increased Ca^2+^ in the chloroplast and upregulates its (p)ppGpp synthase activity (Fig. 7). *CRSH* expression shows a diurnal rhythm with an expression peak at dusk (Mizusawa et al. 2008), which may contribute to the rapid and efficient accumulation of ppGpp during nighttime under natural conditions. The accumulated ppGpp is rapidly degraded by RSH1, RSH2, and RSH3; this regulation may be necessary to avoid the over-suppression of photosynthetic activity and chloroplast metabolism. Once regulation was impaired, plant growth was significantly influenced under photoautotrophic growth conditions. Notably, *crsh-2* still showed a small increase in ppGpp 10 min after the onset of dark treatment, indicating that the increment is not due to the upregulation of CRSH activity; the elucidation of this mechanism will be required in future studies.

**Fig. 7.**
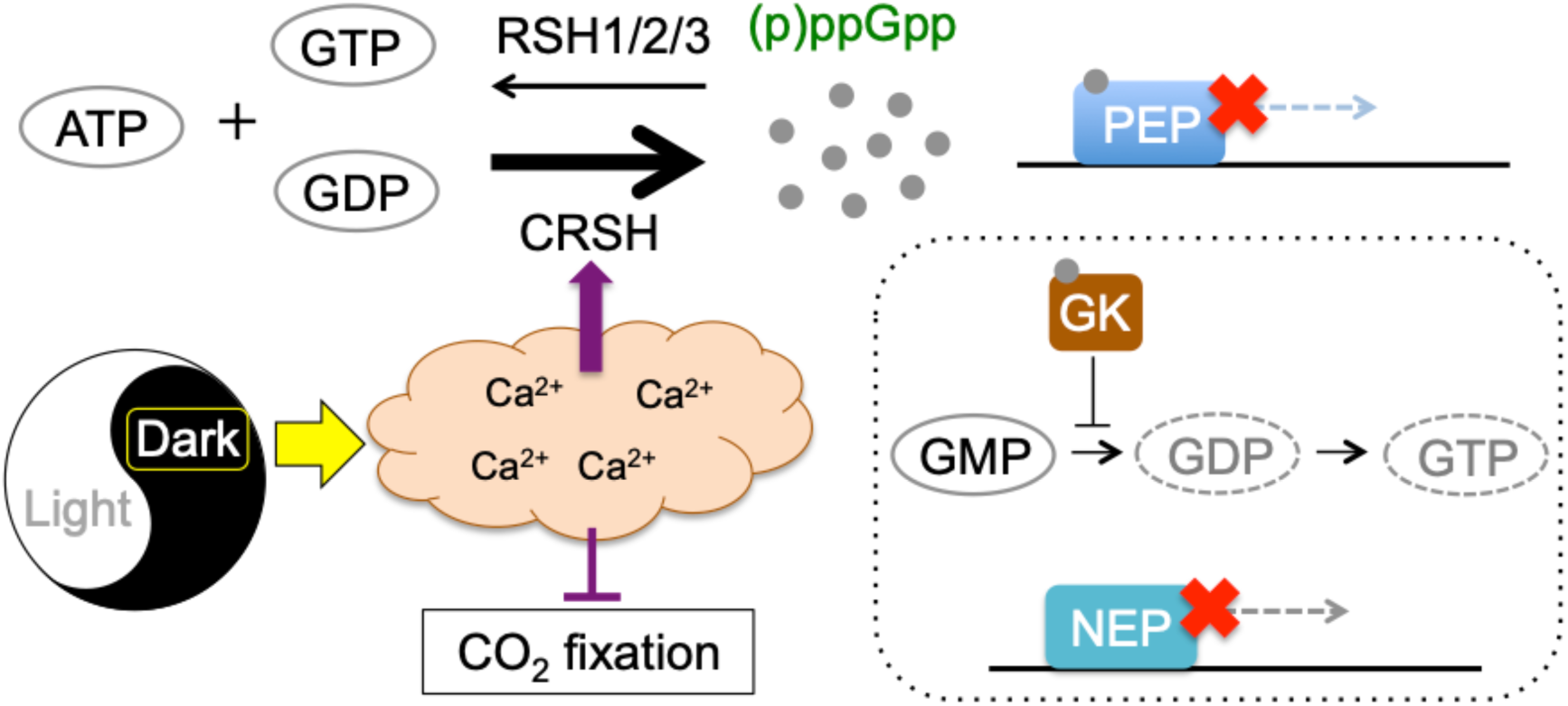
Schematic of dark-induced ppGpp synthesis and regulation of photosynthesis. Upon light-to-dark transition, the plastidial Ca^2+^ concentration is increased, which results in the upregulation of CRSH-dependent (p)ppGpp synthesis and the downregulation of CO_2_ fixation through the inhibition of Calvin–Benson cycle enzymes. Synthesized (p)ppGpp directly inhibits the transcriptional activity of PEP. (p)ppGpp potentially inhibits plastidial guanylate kinase (GK) activity, which results in indirect regulation of transcription by NEP.

ppGpp synthesis by CRSH under dark conditions accompanies the inhibition of PEP-dependent *psbD* transcription in the WT and *rsh2rsh3*. ppGpp directly interacts with PEP *in vitro* to suppress its transcriptional activity (Sato et al. 2009), suggesting that ppGpp, which accumulates upon light-to-dark transition, binds to PEP and inhibits its activity. The mRNA levels of *atpB*, which is transcribed by PEP and NEP, were decreased upon the transition to darkness in *rsh2rsh3*, and super-accumulation of ppGpp was seen, supporting the hypothesis that ppGpp negatively regulates PEP activity. However, we found that transcripts of another PEP-dependent gene, *rbcL*, were not downregulated upon the shift into darkness. The transcript levels of *psbD* and *atpB* in the light are significantly higher than those in the dark; however, the transcript levels of *rbcL* are constant, irrespective of the light conditions (Nakamura et al. 2003). These observations suggest that the significance of the ppGpp-dependent inhibition of PEP activity is variable in each promoter.

Our and another groups previously reported that RSH3-overexpression lines, which constantly accumulate excess ppGpp, suppress the accumulation of transcripts of PEP- and NEP-dependent genes (Maekawa et al. 2015; Sugliani et al. 2016), suggesting that ppGpp also regulates the transcription of NEP-dependent genes by inhibiting guanylate kinase activity and thus reducing the GTP pools necessary for mRNA synthesis (Fig. 7). As direct transcriptional repression of ppGpp may be faster than indirect transcriptional repression through the reduction of GTP pools, indirect transcriptional repression of NEP-dependent genes could not be detected during the timescale of this experiment. Notably, mRNA of *clpP*, which is transcribed by NEP, is transiently increased in *rsh2rsh3* upon the shift to dark conditions. Messenger RNA levels of *clpP* are significantly (>fivefold) higher in the light than in the dark; however, those of *accD* are constant, irrespective of the light conditions (Nakamura et al. 2003). These findings suggest that RSH2 and/or RSH3 are involved in the control of NEP activity and/or the post-transcriptional mRNA degradation of each chloroplast gene, although the mechanism by which this case is carried out remains unclear.

Transient increments in ppGpp levels upon light-to-dark transition have also been reported in the cyanobacterium *Synechococcus elongatus* (Hood et al. 2016; Puszynska and O’Shea 2017), which indicates that the ppGpp-dependent stringent response is universally required to adapt photosynthetic processes to dark conditions in oxygenic phototrophs. Given that cyanobacteria do not contain CRSH homologs (Ito et al. 2017), different strategies must be used to upregulate ppGpp synthesis in the dark in a Ca^2+^-independent manner. Darkness-induced ppGpp accumulation in cyanobacteria may be caused by the posing of the redox signal derived from photosynthetic electron transfer (Hood et al. 2016). Notably, in cyanobacteria, accumulated (p)ppGpp can regulate the expression of all photosynthesis-related genes at transcriptional and translational levels because the genes are located on a single chromosome. However, plants utilize the rise in plastidial Ca^2+^ concentrations and Ca^2+^-dependent ppGpp synthesis to downregulate photosynthetic light and dark reactions in a coordinated manner. Elevated levels of (p)ppGpp and Ca^2+^ negatively control the PEP-dependent transcription of plastid genes, as determined in this study, and Calvin–Benson cycle activities (Portis and Heldt 1976; Charles and Halliwell 1980; Hertig and Wolosiuk 1980; Rocha et al. 2014), respectively. Ca^2+^-dependent (p)ppGpp synthesis may have been required after the endosymbiosis of the ancestral cyanobacterium, because most enzymes that are required for CO_2_ fixation are encoded in the nucleus of plant cells and must be regulated in chloroplasts at the post-translational level.

Although flg22 treatment increases plastidial Ca^2+^ concentrations (Nomura and Shiina 2014), no significant differences were observed in the PAMP regulation of nuclear gene expression in the WT and *crsh-2*, which suggests that the CRSH-dependent stringent response does not interact with PAMP-induced signaling. However, the accumulation of ppGpp has previously been found to affect the susceptibility of *Arabidopsis* to *Turnip mosaic virus*, suggesting that (p)ppGpp metabolism is of physiological importance in plant defense response (Abdelkefi et al. 2018). Furthermore, we found that *WRKY33* transcripts increased upon the transition to darkness and the increment in these transcripts in *rsh2rsh3* 0.5 h after the start of dark incubation was ~2-fold higher than that in the WT, which supports the hypothesis that the ppGpp-dependent plastidial stringent response is involved in plant–pathogen interactions. Perhaps, however, the increase in Ca^2+^ following treatment with 1 µM flg22 is insufficiently high to upregulate CRSH activity to increase plastidial (p)ppGpp levels. The maximum concentration of Ca^2+^ is ~0.1 µM after 1 µM flg22 treatment, which is lower than the value after light-to-dark transition (~0.2 µM) (Nomura et al. 2012), which supports this hypothesis. Future studies should test whether plastidial Ca^2+^ concentrations are increased by the application of other PAMPs; under such conditions, CRSH activity should be upregulated.

In summary, CRSH synthesizes (p)ppGpp upon light-to-dark transition when the Ca^2+^ concentration in chloroplasts increases to a certain level, which is important for plants to adapt plastidial gene expression and metabolism to dark conditions. The *rsh2rsh3* and *rsh1rsh2rsh3* mutants showed significantly higher ppGpp accumulation and the raised levels were maintained longer than those in the WT, indicating that the hydrolase activities of RSH1, RSH2, and RSH3 were required for the adequate maintenance of ppGpp levels in the dark. Further work to determine other conditions in which CRSH-dependent ppGpp synthesis is upregulated will be important in revealing the physiological importance of CRSH function and the relationship between the plastidial stringent response and intracellular Ca^2+^ signaling.

## Materials and Methods

### Plant materials and growth conditions

Plants (Columbia ecotype of *A. thaliana*) were grown on a half concentration of Murashige and Skoog (1/2 MS) medium containing 0.8% agar at 23 °C. Sucrose was added to 1% (w/v) when appropriate. Light conditions are indicated in each experimental procedure. The *rsh2*-*rsh3* double mutant was described previously (Maekawa et al. 2015). The *rsh1* (GABI_206D_08) and *crsh-1* (SAIL_1295_C04) mutants were obtained from the Arabidopsis Biological Resource Center. The *rsh1-rsh2-rsh3* triple mutant was obtained by crossing the *rsh1* mutant with the *rsh2rsh3* mutant. The *crsh*-null mutant, namely, *crsh-2*, was constructed using the CRISPR/Cas9-based method, as described previously (Fauser et al. 2014). We designed a protospacer sequence in the first exon of *CRSH* (83–102 nucleotides from the translation start site), and the protospacer oligonucleotides were annealed and cloned into the plasmid pEn-Chimera based on the protocol (Fauser et al. 2014). After checking the correct sequences of the inserted DNA, it was further cloned into pDe-CAS9 by using Gateway LR Clonase (ThermoFisher). The resultant plasmid was introduced into the *Arabidopsis* WT through the *Agrobacterium*-based standard flower-dip method to induce the double-strand break at the protospacer position. Using gene-specific primers (Table S1), homozygous mutations of *rsh1, rsh2, rsh3*, and/or *crsh* were confirmed by PCR followed by sequencing. RNA was isolated using the SV Total RNA Isolation System (Promega) to check for the presence of the *CRSH* transcript in *crsh-1* (Fig. 2C). The first-strand cDNA, synthesized using the PrimeScript RT regent Kit (TaKaRa), and the genome DNA isolated from the WT were used as templates for RT-PCR with gene-specific primers (Table S1).

The *crsh-2* complementing line was constructed as follows. Previously, we cloned the *CRSH* cDNA (without the stop codon) into the plasmid pDONR/Zeo (Invitrogen) (Masuda et al. 2008). Using the plasmid as a template, five point (but silent) mutations were introduced in the protospacer sequence used to construct *crsh-2* to ensure that the introduced *CRSH* was not recognized by the guide RNA of the CRISPR/Cas9 system. Forward and reverse primers were designed to contain the five mutations (Table S1); the CRSH cDNA, together with the plasmid backbone, was amplified by PCR. The In-Fusion cloning reagent (Clonetech) was used to circularize the amplified fragment. After checking the correct sequence of the plasmid DNA, the insert DNA was cloned into the pGWB2 vector (Nakagawa et al. 2007) by using Gateway LR Clonase (ThermoFisher). Because the cloned *CRSH* did not contain the stop codon, the resulting pGWB2-base plasmid expressed the CRSH fused with an ~1.5 kDa short peptide (YPAFLYKVVDNSA) at the C-terminus; the recombinant protein was designated as CRSH*. The resulting plasmid was introduced into the *crsh-2* mutant by using the *Agrobacterium*-based standard flower-dip method. The expression of CRSH* by the 35S promoter in the complementing line was confirmed by western blot analysis using the anti-CRSH antibody (Supplementary Fig. S2A).

### Quantitative mRNA analysis

To measure the mRNA levels of *accD, clpP, atpB, rbcL, psbD*, and *WRKY33* upon transition to darkness, plants were grown on agar-solidified 1/2 MS medium under continuous light (40 µmol photons m^−2^ s^−1^) for 14 days. After harvesting shoots of the plants (0 h), the remaining plants were transferred to the dark and shoots were harvested after 0.5 and 1 h incubation. Total RNA was isolated using the SV Total RNA Isolation System (Promega). The first-strand cDNA, synthesized using the PrimeScript RT regent Kit (TaKaRa), was the subsequent template. qRT-PCR was performed using SYBR^®^ Premix EX Taq™ and Thermal Cycler Dice^®^ (TaKaRa). Plastid genes, and *WRKY33*-specific primers used for qRT-PCR were as described (Sano et al. 2014; Maekawa et al. 2015).

For the quantification of mRNA levels of *WRKY33* and *WRKY46* after flg22 treatment, plants were grown on agar-solidified 1/2 MS medium under 16 h light (100 µmol photons m^−2^ s^−1^)/16 h dark cycles. Fourteen-day-old plants were treated with 1 µM flg22 (synthesized by Biologica, Nagoya) for 30 min, as described previously (Sano et al. 2014). Total RNA isolation, cDNA synthesis, and gene-specific primers used for qRT-PCR analysis were as described (Sano et al. 2014).

### Quantification of ppGpp

Plants were grown on 0.8% agar-solidified 1/2 MS medium under continuous light (40 µmol photons m^−2^ s^−1^). Fourteen-day-old plants were transferred to the dark and ~0.1 g shoots were harvested at each of the time points indicated (Fig. 3). Extraction and quantification of ppGpp were as described previously (Ihara et al. 2015).

### Western blot analysis

Protein content was measured using the Bradford assay Kit (Bio Lab). Total proteins (6 µg each) were applied to an SDS-PAGE gel and electroblotted onto a polyvinylidene difluoride membrane (GE-Healthcare). The membrane was incubated with the CRSH-specific antibody (Masuda et al. 2008). Immunoreactive proteins were detected using the Alkaline Phosphatase Substrate Kit II (Vector Laboratories).

## Acknowledgments

We thank Professor Holger Puchta (Karlsruhe Institute of Technology) for kindly providing vectors for the CRISPR/Cas9-based genome editing and the Arabidopsis Biological Resource Center for providing mutant seeds.

## Funding

This work was supported by MEXT/JSPS KAKENHI (grant number 19H04719) to SM.

## Disclosures

The authors have no conflicts of interest to declare.

## Abbreviations

CRSH: Ca^2+^-activated RSH
flg22: flagellin 22
(p)ppGpp: 5’-di(tri)phosphate 3’-diphosphate
PAMPs: pathogen-associated molecular patterns
RSH: RelA-SpoT homolog
qRT-PCR: quantitative real-time PCR

---

**Table S1.**
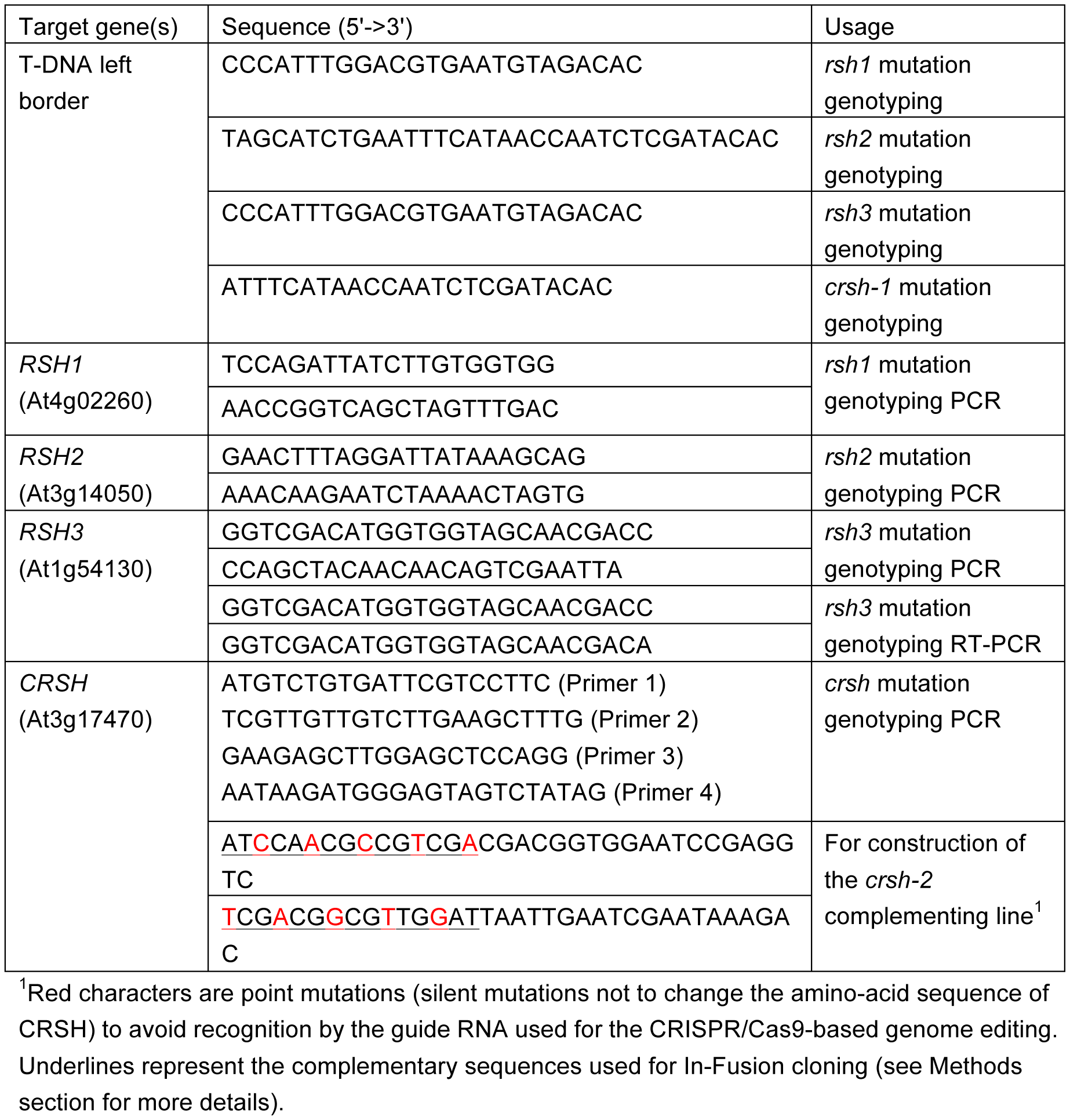
Primers used in this study

**Fig. S1.**
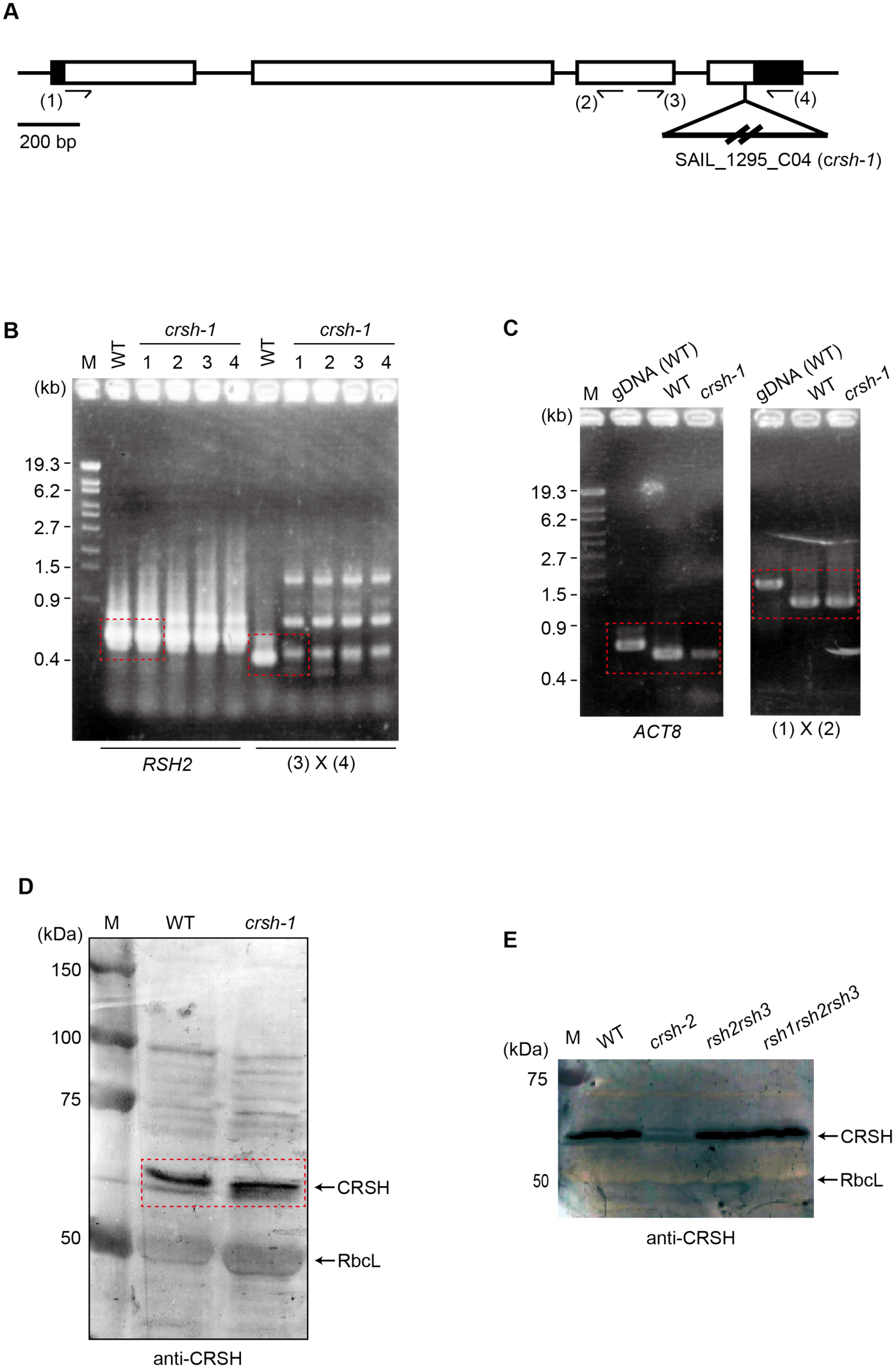
Original photographs and western blot data used for Fig. 1. (A) Schematic of the exon– intron structure of *CRSH* (At3g17470) with a single nucleotide deletion site in *crsh-2* and a T-DNA insertion site in *crsh-1*. (B) PCR analysis reveals homozygous T-DNA insertion in *crsh-1*. Positions of primers used (3 and 4) are indicated in (A). An asterisk indicates a nonspecific band. *RSH2* was amplified as a control for DNA isolation. (C) RT-PCR analysis reveals the presence of *CRSH* mRNA in *crsh-1*. Positions of primers used (1 and 2) are indicated in (A). Genome DNA isolated from WT was used as a control for genome DNA contamination, and *actin8* (ACT8) gene was used as a control for mRNA isolation and reverse transcription. (D) Western blot of proteins isolated from WT and *crsh-1*. (E) Western blot of proteins isolated from WT, *crsh-2, rsh2rsh3*, and *rsh1rsh2rsh3*. Red perpendiculars indicate the positions used for constructing Fig. 1.

**Fig. S2.**
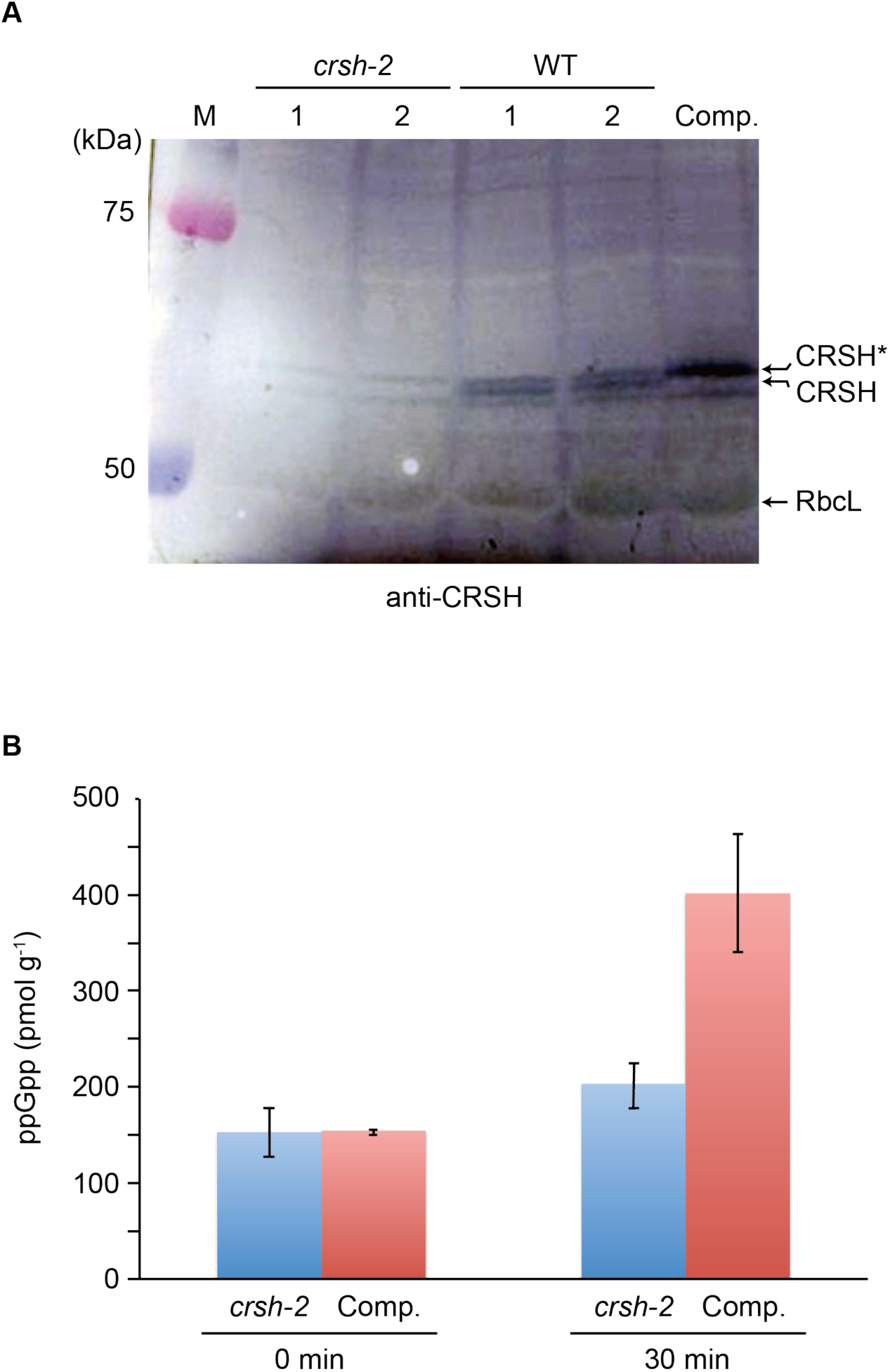
Characterization of *crsh-2* complementing line with 35S-promoter-driven CRSH having additional 13 amino acids (~1.5 kDa) at the C-terminus (CRSH*). (A) Western blot of proteins isolated from WT, *crsh-2*, and *crsh-2* complementing lines expressing CRSH* (Comp.). (B) ppGpp accumulation before (0 min) and after dark incubation for 30 min (30 min) in *crsh-2* and the complementing line (Comp.). Data are means of three biological replicates.

**Fig. S3.**
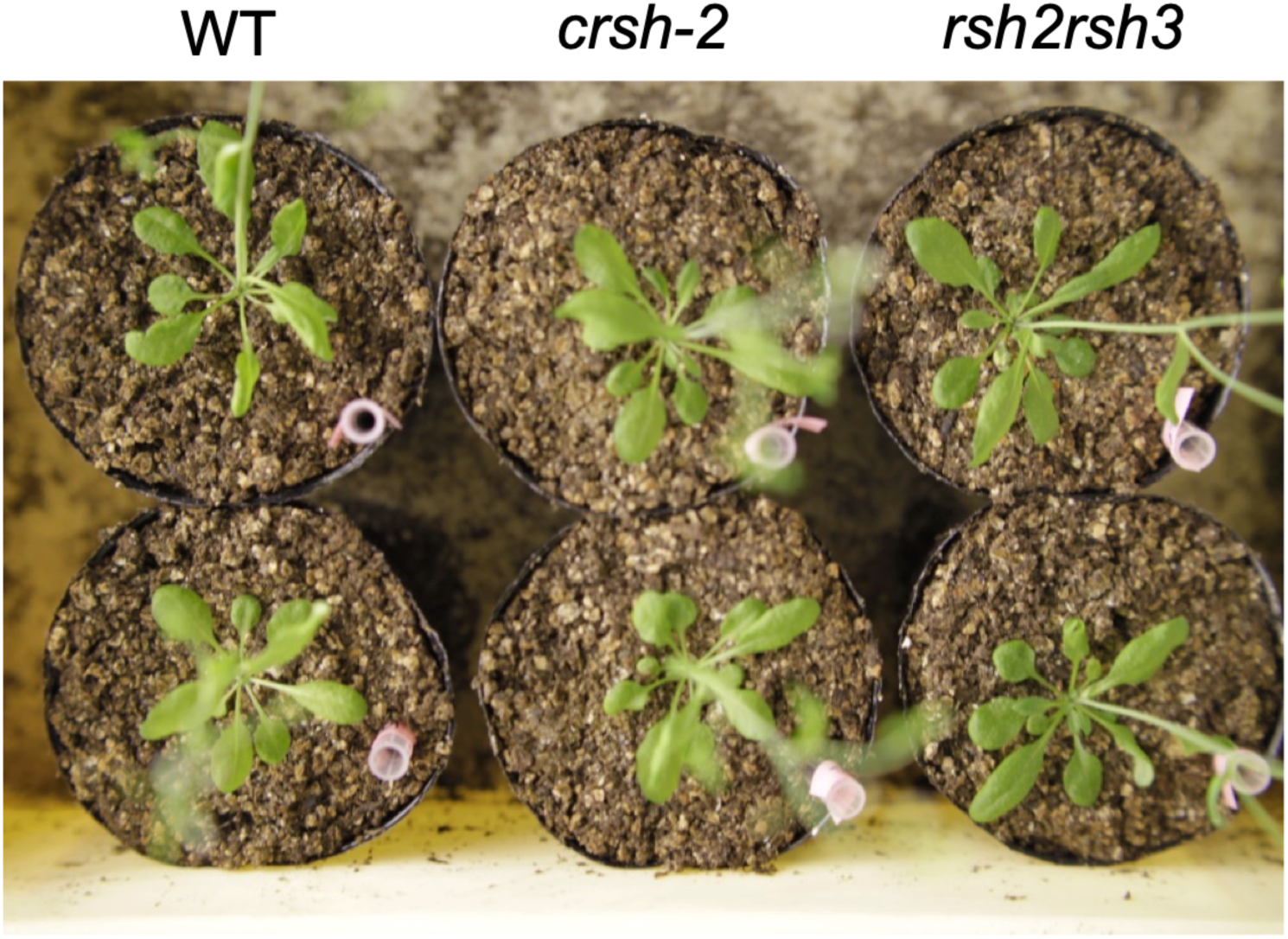
No significant visible phenotype was observed for *crsh-2* and *rsh2rsh3* mutants. Plants were germinated on 1/2 MS plates and grown for 2 weeks. Plants were then transferred to soil and grown for 1 month under continuous-light conditions.

